# Finding the right words to evaluate research: An empirical appraisal of eLife’s assessment vocabulary

**DOI:** 10.1101/2024.04.30.591844

**Authors:** Tom E. Hardwicke, Sarah R. Schiavone, Beth Clarke, Simine Vazire

## Abstract

Research articles published by the journal eLife are accompanied by short evaluation statements that use phrases from a prescribed vocabulary to evaluate research on two dimensions: importance and strength of support. Intuitively, the prescribed phrases appear to be highly synonymous (e.g., important/valuable, compelling/convincing) and the vocabulary’s ordinal structure may not be obvious to readers. We conducted an online repeated-measures experiment to gauge whether the phrases were interpreted as intended. We also tested an alternative vocabulary with (in our view) a less ambiguous structure. 301 participants with a doctoral or graduate degree used a 0-100% scale to rate the importance and strength of support of hypothetical studies described using phrases from both vocabularies. For the eLife vocabulary, most participants’ implied ranking did not match the intended ranking on both the importance (*n* = 59, 20% matched, 95% confidence interval [15% to 24%]) and strength of support dimensions (*n* = 45, 15% matched [11% to 20%]). By contrast, for the alternative vocabulary, most participants’ implied ranking did match the intended ranking on both the importance (*n* = 188, 62% matched [57% to 68%]) and strength of support dimensions (*n* = 201, 67% matched [62% to 72%]). eLife’s vocabulary tended to produce less consistent between-person interpretations, though the alternative vocabulary still elicited some overlapping interpretations away from the middle of the scale. We speculate that explicit presentation of a vocabulary’s intended ordinal structure could improve interpretation. Overall, these findings suggest that more structured and less ambiguous language can improve communication of research evaluations.

## INTRODUCTION

Peer review is usually a black box — readers only know that a research paper eventually surpassed some ill-defined threshold for publication and rarely see the more nuanced evaluations of the reviewers and editor (Vazire, 2021). A minority of journals challenge this convention by making peer review reports publicly available (Wolfram et al., 2020). One such journal, eLife, also accompanies articles with short evaluation statements (“eLife assessments”) representing the consensus opinions of editors and peer reviewers (Eisen et al., 2022). In 2022, eLife stated that these assessments would use phrases drawn from a common vocabulary (Table 1) to convey their judgments on two evaluative dimensions: (1) “significance”; and (2) “strength of support” (for details see *eLife’s New Model*, 2022). For example, a study may be described as having “landmark” significance and offering “exceptional” strength of support (for a complete example, see Box 1). The phrases are drawn from “widely-used expressions” in prior eLife assessments and the stated goal is to “help convey the views of the editor and the reviewers in a clear and consistent manner” (*eLife’s New Model*, 2022). Here we report a study which assessed whether the language used in eLife assessments is perceived clearly and consistently by potential readers. We also assessed alternative language that may improve communication.

**Table 1.** Phrases and their definitions (italicised) from the eLife vocabulary representing two evaluative dimensions: significance and strength of support. The significance dimension is represented by five phrases and the strength of support dimension is represented by six phrases. In a particular eLife assessment, readers only see one phrase from each of the evaluative dimensions. Phrases are accompanied by eLife definitions, but these are not shown in eLife assessments (though some words from the definitions may be used).

| **eLife vocabulary** |  |
| --- | --- |
| **Significance** | **Strength of support** |
| Landmark: *findings with profound implications that are expected to have widespread influence* | Exceptional: *exemplary use of existing approaches that establish new standards for a field* |
| Fundamental: *findings that substantially advance our understanding of major research questions* | Compelling: *evidence that features methods, data and analyses more rigorous than the current state-of-the-art* |
| Important: *findings that have theoretical or practical implications beyond a single subfield* | Convincing: *appropriate and validated methodology in line with current state-of-the-art* |
| Valuable: *findings that have theoretical or practical implications for a subfield* | Solid: *methods, data and analyses broadly support the claims with only minor weaknesses* |
| Useful: *findings that have focused importance and scope* | Incomplete: *main claims are only partially supported* |
|  | Inadequate: *methods, data and analyses do not support the primary claims* |

Our understanding (based on *eLife’s New Model*, 2022) is that eLife intends the common vocabulary to represent different degrees of each evaluative dimension on an ordinal scale (e.g., “landmark” findings are more significant than “fundamental” findings, and so forth); however, in our view the intended ordering is sometimes ambiguous or counterintuitive. For example, it does not seem obvious to us that an “important” study is necessarily more significant than a “valuable” study, nor does a “compelling” study seem necessarily stronger than a “convincing” study. Additionally, several phrases like “solid” and “useful”, could be broadly interpreted, leading to a mismatch between intended meaning and perceived meaning. The phrases also do not cover the full continuum of measurement and are unbalanced in terms of positive and negative phrases^1^. For example, the “significance” dimension has no negative phrases — the scale endpoints are “landmark” and “useful”. We also note that the definitions provided by eLife do not always map onto gradations of the same construct. For example, the eLife definitions of phrases on the significance dimension suggest that the difference between “useful”, “valuable”, and “important” is a matter of breadth/scope (whether the findings have implications beyond a specific subfield) whereas the difference between “fundamental” and “landmark” is a matter of degree. In short, we are concerned that several aspects of the eLife vocabulary may undermine communication of research evaluations to readers.

In Table 2, we outline an alternative vocabulary that is intended to overcome these potential issues with the eLife vocabulary. Phrases in the alternative vocabulary explicitly state the relevant evaluative dimension (e.g., “support”) along with a modifying adjective that unambiguously represents degree^2^ (e.g., “very low”). The alternative vocabulary is intended to cover the full continuum of measurement and be balanced in terms of positive and negative phrases. We have also renamed “significance” to “importance” to avoid any confusion with statistical significance. We hope that these features will facilitate alignment of readers’ interpretations with the intended interpretations, improving the efficiency and accuracy of communication.

**Table 2.** Phrases from the alternative vocabulary representing two evaluative dimensions: importance and strength of support. Each dimension is represented by five phrases.

| **Alternative vocabulary** |  |
| --- | --- |
| **Importance** | **Strength of support** |
| Very high importance | Very strong support |
| High importance | Strong support |
| Moderate importance | Moderate support |
| Low importance | Weak support |
| Very low importance | Very weak support |

### Box 1. A complete example of an eLife assessment. This particular example uses the phrase “important”, to convey the study’s significance, and the phrase “compelling”, to convey the study’s strength of support.

“The overarching question of the manuscript is important and the findings inform the patterns and mechanisms of phage-mediated bacterial competition, with implications for microbial evolution and antimicrobial resistance. The strength of the evidence in the manuscript is compelling, with a huge amount of data and very interesting observations. The conclusions are well supported by the data. This manuscript provides a new co-evolutionary perspective on competition between lysogenic and phage-susceptible bacteria, that will inform new studies and sharpen our understanding of phage-mediated bacterial co-evolution.” (Rendueles et al., 2023).

The utility of eLife assessments will depend (in part) on whether readers interpret the common vocabulary in the manner that eLife intends. Mismatches between eLife’s intentions and readers’ perceptions could lead to inefficient or inaccurate communication. In this study, we empirically evaluated how the eLife vocabulary (Table 1) is interpreted and assessed whether an alternative vocabulary (Table 2) elicited more desirable interpretations. Our goal was not to disparage eLife’s progressive efforts, but to make a constructive contribution towards a more transparent and informative peer review process. We hope that a vocabulary with good empirical performance will be more attractive and useful to other journals considering adopting eLife’s approach.

Our study is modelled on prior studies that report considerable individual differences in people’s interpretation of probabilistic phrases (Budescu et al., 2014; Budescu & Wallsten, 1985; Lichtenstein & Newman, 1967; Reagan et al., 1989; Theil, 2002; Wallsten et al., 1986; Willems et al., 2020). In a prototypical study of this kind, participants are shown a probabilistic statement like “It will probably rain tomorrow” and asked to indicate the likelihood of rain on a scale from 0-100%. Analogously, in our study participants read statements describing hypothetical scientific studies using phrases drawn from the eLife vocabulary or the alternative vocabulary and were asked to rate the study’s significance/importance or strength of support on a scale from 0-100. We used these responses to gauge the extent to which people’s interpretations of the vocabulary were consistent with each other and consistent with the intended rank order.

### Research aims

Our overarching goal was to identify clear language for conveying evaluations of scientific papers. We hope that this will make it easier for other journals/platforms to follow in eLife’s footsteps and move towards more transparent and informative peer review.

With this overall goal in mind, we had three specific research aims:

- Aim One. To what extent do people share similar interpretations of phrases used to describe scientific research?
- Aim Two. To what extent do people’s (implied) ranking of phrases used to describe scientific research align with (a) each other; and (b) with the intended ranking?
- Aim Three. To what extent do different phrases used to describe scientific research elicit overlapping interpretations and do those interpretations imply broad coverage of the underlying measurement scale?

## METHODS

Our methods adhered to our preregistered plan (https://doi.org/10.17605/OSF.IO/MKBTP) with one minor deviation: our target sample size was 300, but we accidentally recruited an additional participant, so the actual sample size was 301.

### Design

We conducted an experiment with a repeated-measures design. Participants were shown short statements that described hypothetical scientific studies in terms of their significance/importance or strength of support using phrases drawn from the eLife vocabulary (Table 1) and from the alternative vocabulary (Table 2). The statements were organised into four blocks based on vocabulary and evaluative dimension; specifically, block one: *eLife-significance* (5 statements), block two: *eLife-support* (6 statements), block three: *alternative-importance* (5 statements), block four: *alternative-support* (5 statements). Each participant saw all 21 phrases and responded using a 0-100% slider scale to indicate their belief about each hypothetical study’s significance/importance or strength of support.

### Materials

There were 21 statements that described hypothetical scientific studies using one of the 21 phrases included in the two vocabularies (Table 1). Statements referred either to a study’s strength of support (e.g., Figure 1) or a study’s significance/importance (e.g., Supplementary Figure A1). For the alternative vocabulary, we used the term “importance” rather than “significance”. To ensure the statements were grammatically accurate, it was necessary to use slightly different phrasing when communicating significance with the eLife vocabulary (“This is an [phrase] study”) compared to communicating importance with the alternative vocabulary (“This study has [phrase] importance”; e.g., Supplementary Figure A2).

**Figure 1.**
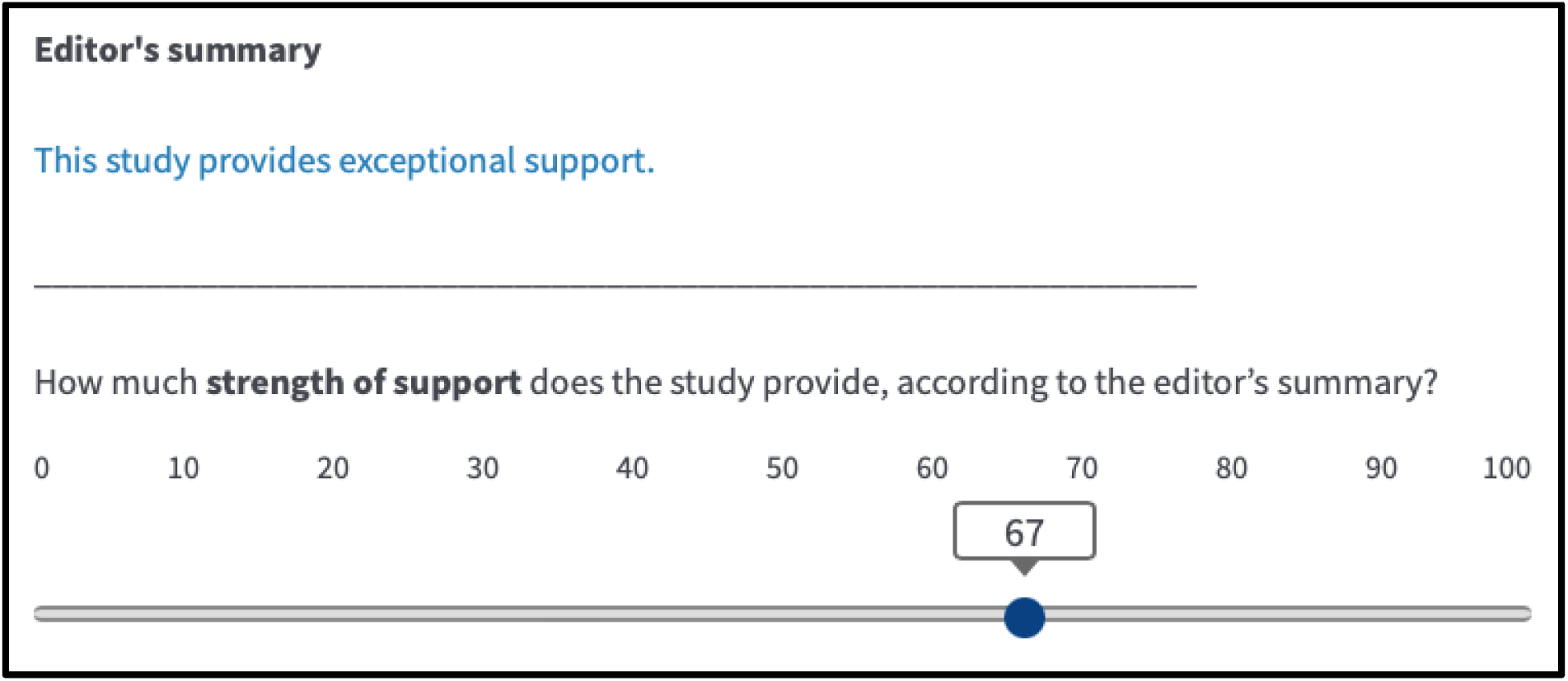
An example summary statement referring to a study’s strength of support and the corresponding response scale with an arbitrary response shown.

**Figure 1:**
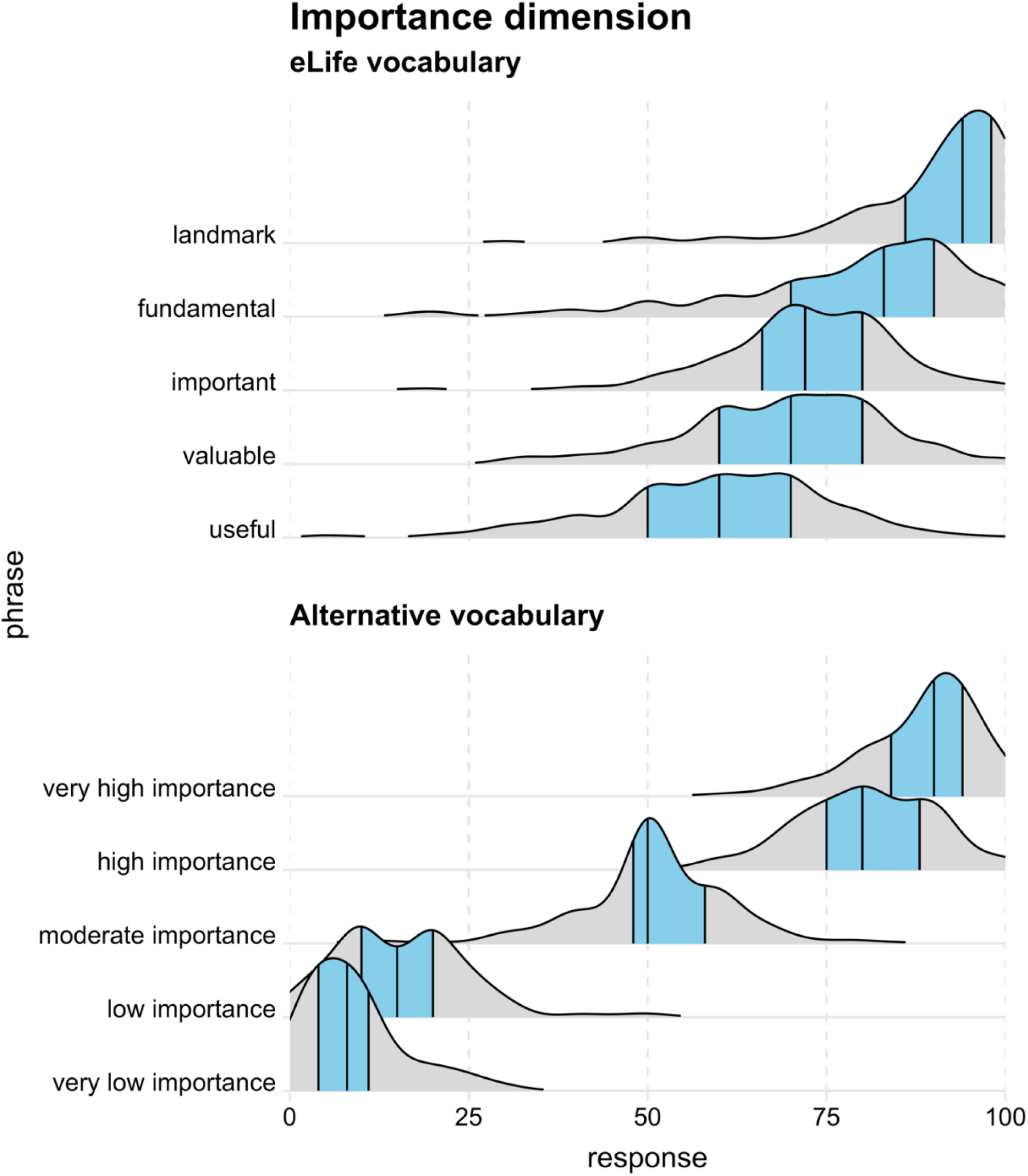
Responses to each phrase on the importance/significance dimension as kernel density distributions with the 25th, 50th (i.e., median), and 75th quantiles represented by black vertical lines and the 25th-75th quantile region (i.e., interquartile range) highlighted in blue.

Additionally, there was one attention check statement (Supplementary Figure A3), a question asking participants to confirm their highest completed education level (options: Undergraduate degree (BA/BSc/other)/Graduate degree (MA/MSc/MPhil/other)/Doctorate degree (PhD/other)/Other), and a question asking participants the broad subject area of their highest completed education level (options: Arts & Humanities/Life Sciences and Biomedicine/Physical Sciences/Social Sciences/Other). The veridical materials are available at https://osf.io/jpgxe/.

### Sample

#### Sample source

Participants were recruited from the online participant recruitment platform Prolific (https://www.prolific.co/). As of 23^rd^ August 2023, the platform had 123,064 members. Demographic information about Prolific members is provided in Supplementary Information H.

#### Sample size

As data collection unfolded, we intermittently checked how many participants had met the inclusion criteria, aiming to stop data collection when we had eligible data for our target sample size of 300 participants. Ultimately, 461 participants responded to the survey. Of these 461 participants, 156 participants failed the attention check and 12 participants took longer than 30 minutes to complete the study and were therefore excluded. No participants failed to respond to all 21 statements or completed the study too quickly (< 5 minutes). We applied these exclusion criteria one-by-one which removed data from 160 participants and retained eligible data from 301 participants (we unintentionally recruited one additional participant).

#### Sample size justification

The target sample size of 300 was based on our resource constraints and expectations about statistical power and precision (see Supplementary Information B).

#### Inclusion criteria

Participants had to have a ≥ 95% approval rate for prior participation on the recruitment platform (Prolific). Additionally, Prolific pre-screening questions were used to ensure that the study was only available to participants who reported that they speak fluent English, were aged between 18-70 years, and had completed a doctorate degree (PhD/other).

### Procedure

1. Data collection and recruitment via the Prolific platform began on September 13^th^ 2023 and was completed on September 14^th^ 2023.
2. After responding to the study advert (https://osf.io/a25vq), participants read an information sheet (https://osf.io/39vay), and provided consent (https://osf.io/xdar7). During this process, they were told that the study seeks to understand “how people perceive words used to describe scientific studies so we can improve communication of research to the general public.”
3. Participants completed the task remotely online via the Qualtrics platform. Before starting the main task, they read a set of instructions and responded to a practice statement (Supplementary Information C).
4. For the main task, statements were presented sequentially, and participants responded to them in their own time. The order of presentation was randomized, both between and within the four blocks of statements. After each statement, there was a 15 second filler task during which participants were asked to complete as many multiplication problems (e.g., 5 x 7 = ?) as they could from a list of 10. The multiplication problems were randomly generated every time they appeared using the Qualtrics software. Only numbers between 1 and 15 were used to ensure that most of the problems were relatively straightforward to solve. A single ‘attention check’ statement (Supplementary Figure A3) appeared after all four blocks had been completed.
5. Participants were required to respond to each statement before they could continue to the next statement. The response slider could be readjusted as desired until the ‘next’ button was pressed, after which participants could not return to or edit prior responses.
6. After responding to all 21 statements and the attention check, participants were shown a debriefing document (https://osf.io/a9gve).

## RESULTS

All analyses adhered to our preregistered plan (https://doi.org/10.17605/OSF.IO/MKBTP). Numbers in square brackets represent 95% confidence intervals computed with the Sison-Glaz method (Sison & Glaz, 1995) for multinomial proportions or bootstrapped with the percentile method (Rousselet et al., 2021) for percentile estimates.

### Participant characteristics

Participants stated that their highest completed education level was either a doctorate degree (*n* = 287) or graduate degree (*n* = 14)^3^. Participants reported that the subject areas that most closely represented their degrees were Life Sciences and Biomedicine (*n* = 97), Social Sciences (*n* = 77), Physical Sciences (*n* = 57), Arts & Humanities (*n* = 37) and various “other” disciplines (*n* = 33).

### Response distributions

The distribution of participants’ responses to each phrase is shown in Figure 1 (importance/significance dimension) and Figure 2 (strength of support dimension). These ‘ridgeline’ plots (Wilke, 2019) are kernel density distributions which represent the relative probability of observing different responses (akin to a smoothed histogram).

**Figure 2:**
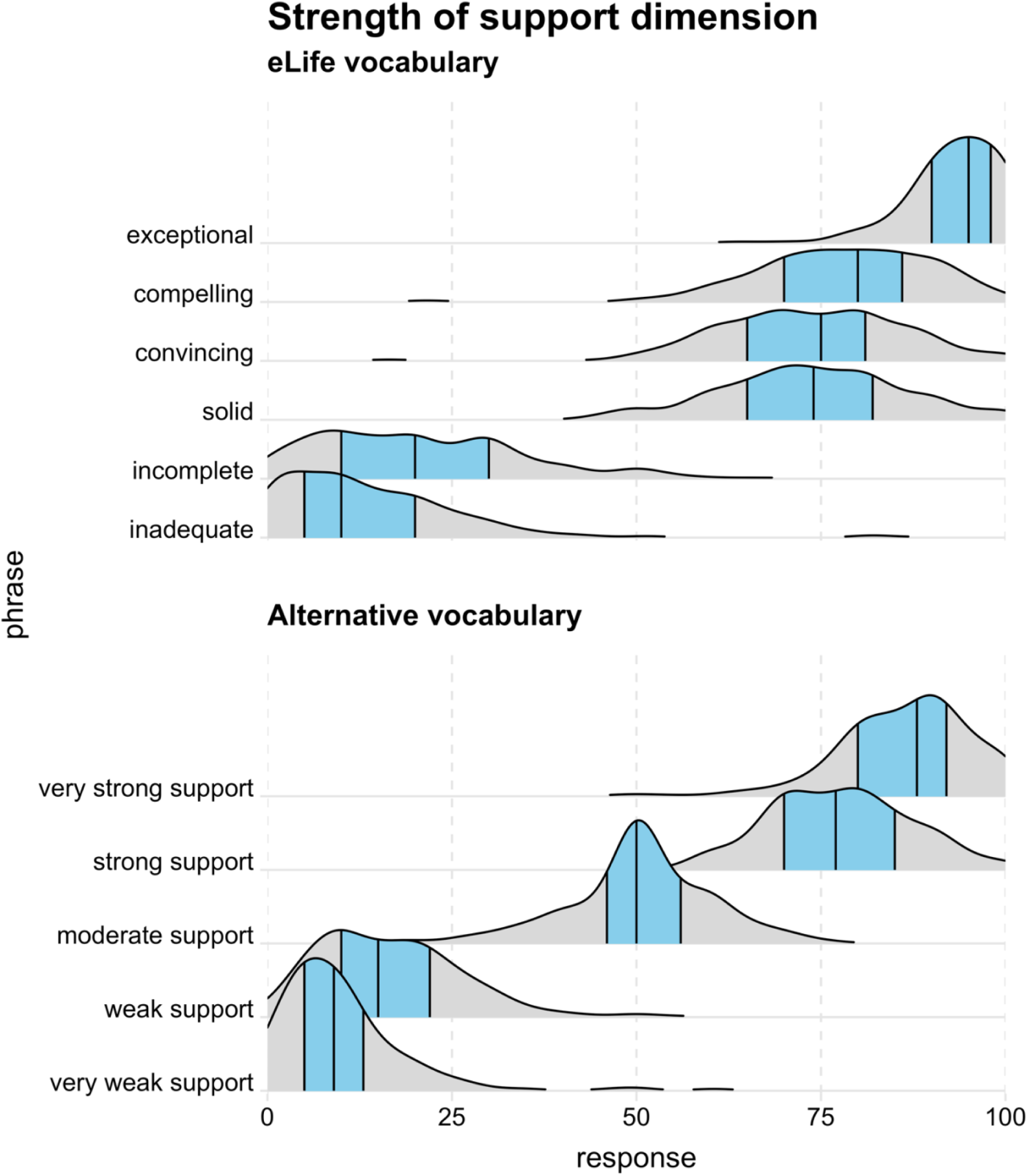
Responses to each phrase on the strength of support dimension as kernel density distributions with the 25th, 50th (i.e., median), and 75th quantiles represented by black vertical lines and the 25th-75th quantile region (i.e., interquartile range) highlighted in blue.

Tables 3 and Table 4 show the 25th, 50th (i.e., median), and 75th percentiles of responses for each phrase (as represented by the black vertical lines in Figure 1 and Figure 2). The tables include 95% confidence intervals only for medians to make them easier to read; however, confidence intervals for all percentile estimates are available in Supplementary Table E1 and Supplementary Table E2.

**Table 3.** Percentile estimates for participant responses to phrases on the significance/importance dimension for eLife and alternative vocabularies.

| **Importance dimension** |  |  |  |
| --- | --- | --- | --- |
| **Phrase** | **Percentile** |  |  |
|  | **25th** | **Median [CI]** | **75th** |
| **eLife vocabulary** |  |  |  |
| Landmark | 86 | 94 [92,95] | 98 |
| Fundamental | 70 | 83 [81,85] | 90 |
| Important | 66 | 72 [71,75] | 80 |
| Valuable | 60 | 70 [70,71] | 80 |
| Useful | 50 | 60 [60,63] | 70 |
| **Alternative vocabulary** |  |  |  |
| Very high importance | 84 | 90 [90,91] | 94 |
| High importance | 75 | 80 [80,81] | 88 |
| Moderate importance | 48 | 50 [50,51] | 58 |
| Low importance | 10 | 15 [13,18] | 20 |
| Very low importance | 4 | 8 [6,9] | 11 |

**Table note:** CI: 95% confidence intervals.

**Table 4.**
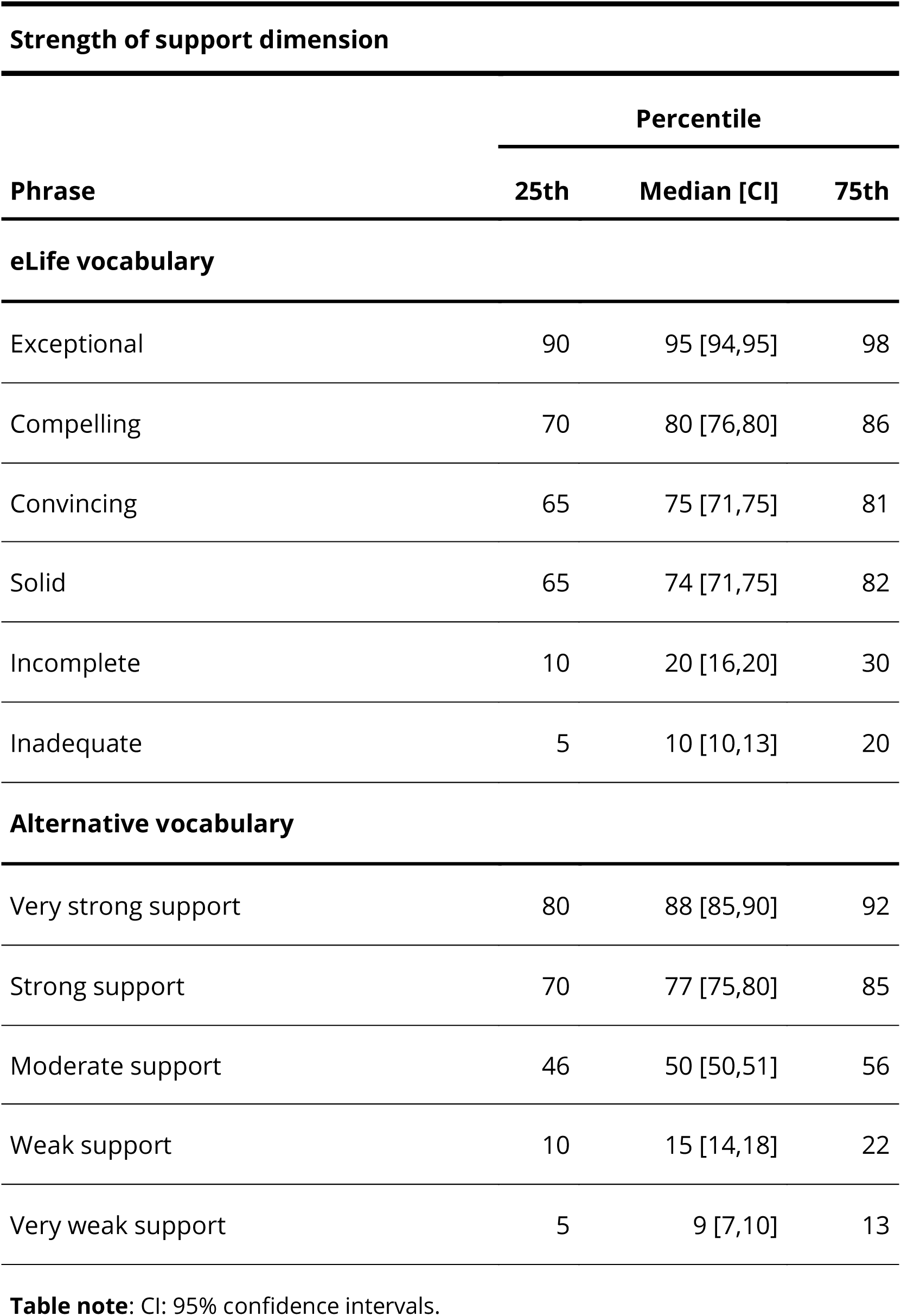
Percentile estimates for participant responses to phrases on the strength of support dimension for eLife and alternative vocabularies.

### Implied ranking of evaluative phrases

#### Do participants’ implied rankings match the intended rankings?

Although participants rated each statement separately on a continuous scale, these responses also imply an overall ranking of the phrases (in order of significance/importance or strength of support). Ideally, an evaluative vocabulary elicits implied rankings that are both consistent among participants and consistent with the intended ranking. Figure 3 shows the proportion of participants whose implied ranking matched the intended ranking (i.e., “correct ranking”) for the different evaluative dimensions and vocabularies.

**Figure 3.**
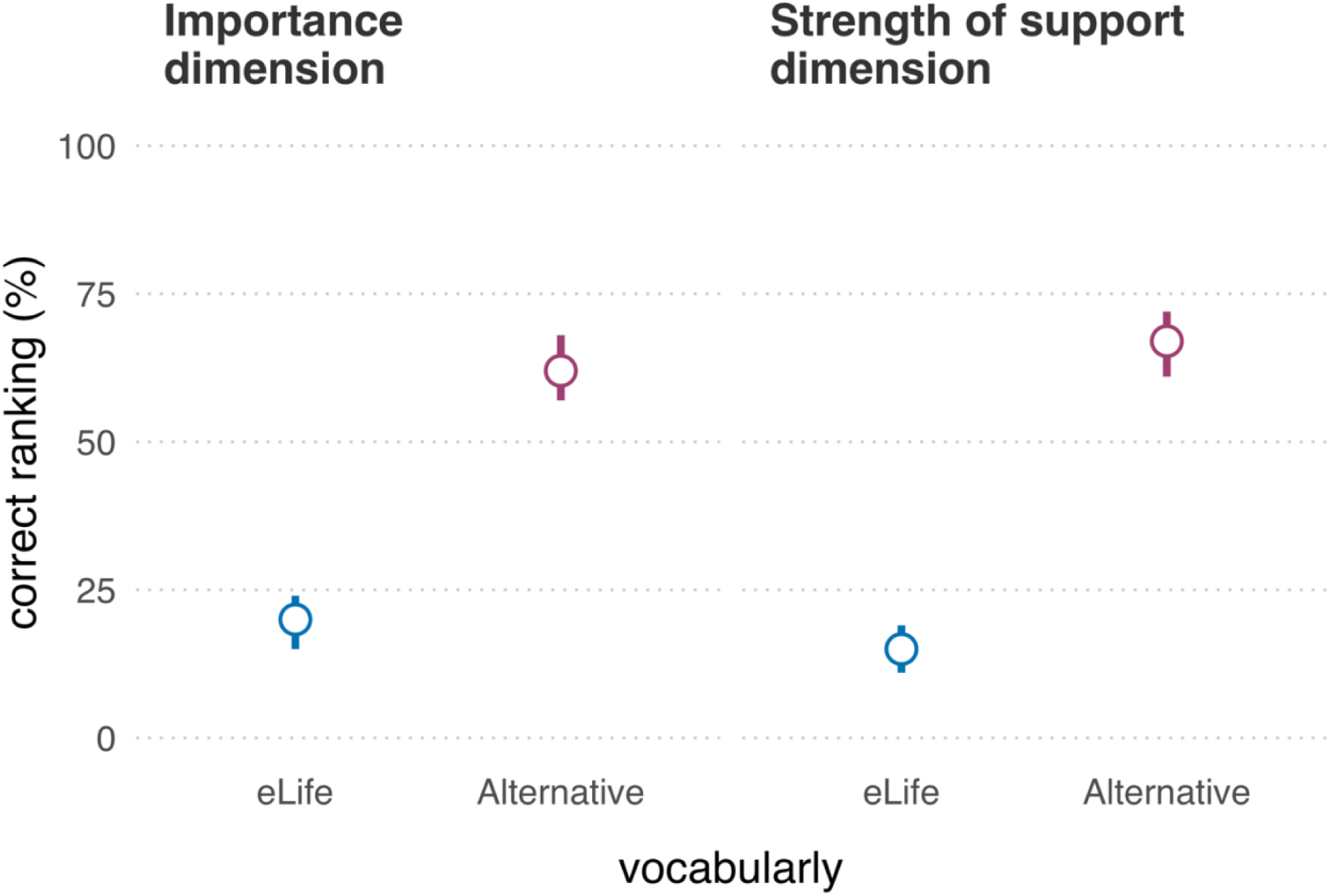
Proportion of participants (N = 301) whose implied ranking matched the intended ranking (i.e., ‘correct ranking’) for both evaluative dimensions and both vocabularies. Error bars represent 95% confidence intervals using the Sison-Glaz method (Sison & Glaz, 1995).

On the significance/importance dimension, 59 (20% [15% to 24%]) participants’ implied rankings of the eLife vocabulary aligned with the intended ranking and 188 (62% [57% to 68%]) participants’ implied rankings of the alternative vocabulary aligned with the intended ranking. We performed an ‘exact’ McNemar test and computed the McNemar odds ratio with Clopper Pearson 95% confidence intervals adjusted with the ‘midp’ method, as recommended by Fagerland et al. (2013). The McNemar test indicated that observing a difference between the vocabularies this large, or larger, is unlikely if the null hypothesis were true (odds ratio = 8.17, 95% CI [5.11,13.69], *p* = 1.34e-26). The intended ranking was the most popular for both vocabularies; however, participants had 55 different implied rankings for the eLife vocabulary and 8 different implied rankings for the alternative vocabulary (for details, see Supplementary Tables F1-F4). Note that these values should be compared with caution, as for the significance/importance dimension, the eLife vocabulary had more (six) phrases than the alternative vocabulary (which had five phrases and therefore fewer possible rankings).

On the strength of support dimension, 45 (15% [11% to 20%]) participants’ ratings of the eLife phrases were in accordance with the intended ranking relative to 201 (67% [62% to 72%]) participants who correctly ranked the alternative vocabulary. A McNemar test indicated that observing a difference between the vocabularies this large, or larger, is unlikely if the null hypothesis were true (odds ratio = 11.4, 95% CI [6.89, 20.01], *p* = 5.73e-35). The intended ranking was the most popular for both vocabularies; though for the eLife vocabulary an unintended ranking that swapped the ordinal positions of “convincing” and “solid” came a close second, reflected in the ratings of 44 (15% [10% to 19%]) participants. Overall, there were 34 different implied rankings for the eLife vocabulary, relative to 10 implied rankings for the alternative vocabulary.

#### Quantifying ranking similarity

Thus far, our analyses have emphasized the binary difference between readers’ implied rankings and eLife’s intended rankings. A complementary analysis quantifies the *degree* of similarity between rankings using Kendall’s tau distance (*K*_d_) — a metric that describes the difference between two lists in terms of the number of adjacent pairwise swaps required to convert one list into the other (Kendall, 1938; van Doorn et al., 2021). The larger the distance, the larger the dissimilarity between the two lists. *K*_d_ ranges from 0 (indicating a complete match) to n(n-1)/2 (where n is the size of one list). Because the eLife strength of support dimension has six phrases and all other dimensions have five phrases, we report the normalised *K*_d_ which ranges from 0 (maximal similarity) to 1 (maximal dissimilarity). Further explanation of *K*_d_ is provided in Supplementary Information G.

Figure 4 illustrates the extent to which participants’ observed rankings deviated from the intended ranking in terms of normalized *K_d_*. This suggests that although deviations from the intended eLife ranking were common, they only tended to be on the order of one or two discordant rank pairs. By contrast, the alternative vocabulary rarely resulted in any deviations, and when it did these were typically only in terms of one discordant rank pair.

**Figure 4.**
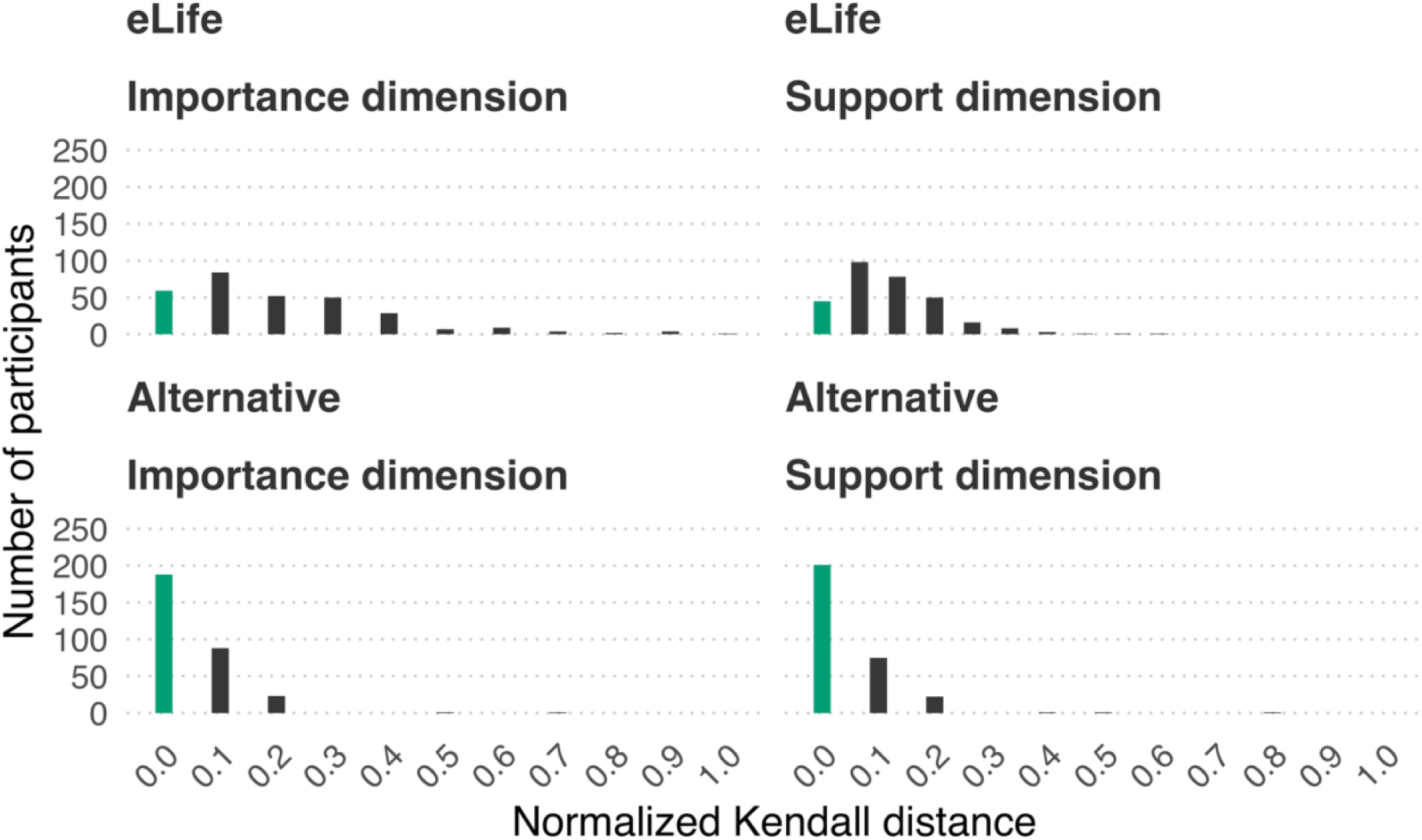
Extent to which participants (N = 301) implied rankings deviated from intended rankings using normalized Kendall’s distance. Zero represents a perfect match (green bar), other values (grey bars) represent increasing dissimilarity from the intended ranking up to 1, which represents maximal dissimilarity.

#### Locus of ranking deviations

So far, we have examined how many participants adhered to the intended ranking (Figure 3) and the extent to which their implied rankings deviated from the intended rankings (Figure 4). However, these approaches do not optimally illustrate *where* the ranking deviations were concentrated (i.e., which phrases were typically being misranked). The heat maps in Figure 5 show the percentage of participants whose implied rankings matched or deviated from the intended ranking at the level of individual phrases. Ideally, a phrase’s observed rank will match its intended rank for 100% of participants. For example, the heat maps show that almost all participants (98%) correctly ranked “moderate importance” and “moderate support” in the alternative vocabulary. The heat maps also reveal phrases that were often misranked with each other, for example: “solid”, “convincing”, and “compelling” in the eLife vocabulary.

**Figure 5.**
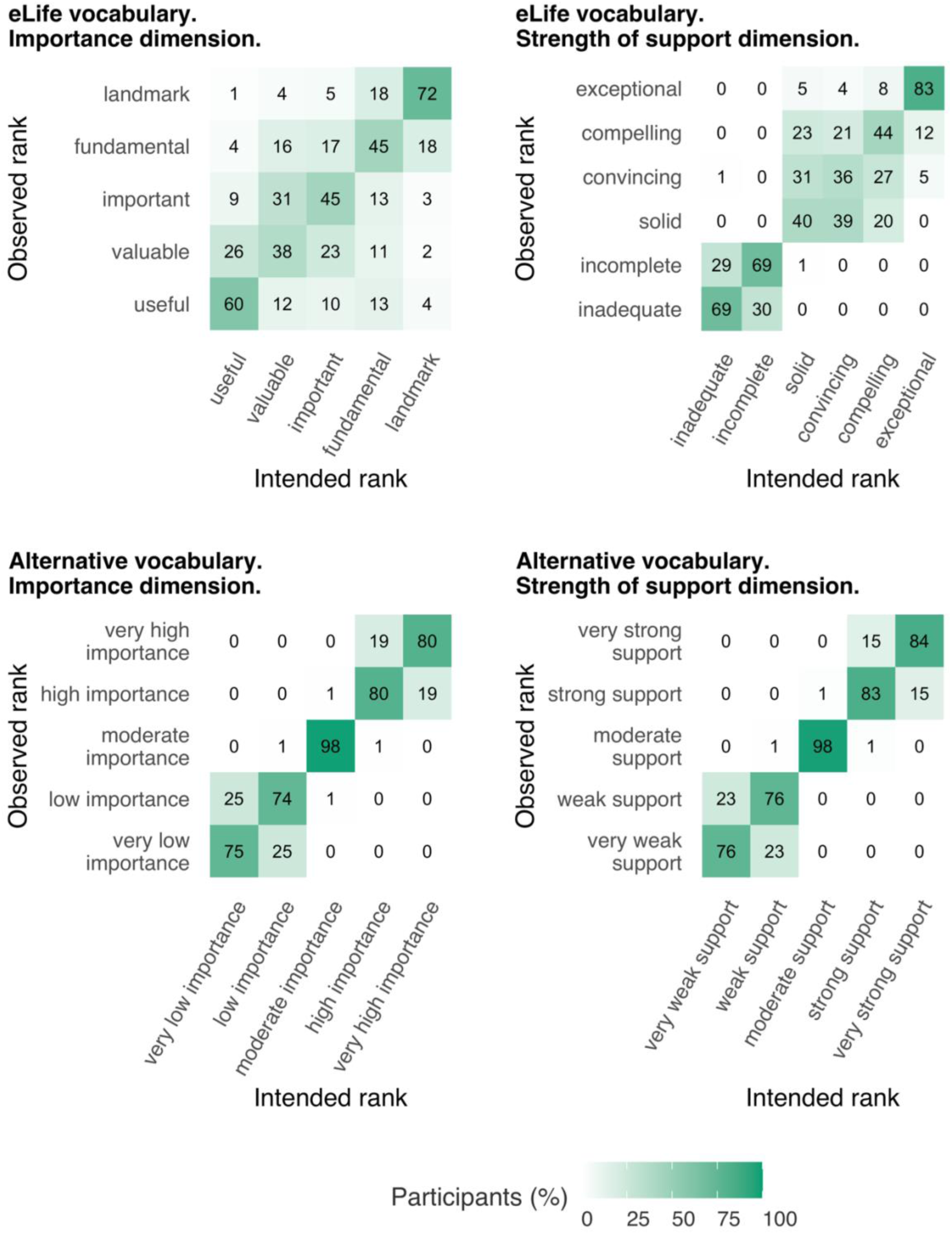
Heat maps showing the percentage of participants (N = 301) whose implied rankings were concordant or discordant with the intended ranking at the level of individual phrases. Darker colours and higher percentages indicate greater concordance between the implied rank and the intended rank of a particular phrases.

## DISCUSSION

Research articles published in eLife are accompanied by evaluation statements that use phrases from a prescribed vocabulary (Table 1) to describe a study’s importance (e.g., “landmark”) and strength of support (e.g., “compelling”). If readers, reviewers, and editors interpret the prescribed vocabulary differently to the intended meaning, or inconsistently with each other, it could lead to miscommunication of research evaluations. In this study, we assessed the extent to which people’s interpretations of the eLife vocabulary are consistent with each other, and consistent with the intended ordinal structure. We also examined whether an alternative vocabulary (Table 2) improved consistency of interpretation.

Overall, the empirical data supported our initial intuitions: while some phrases in the eLife vocabulary were interpreted relatively consistently (e.g., “exceptional” and “landmark”), several phrases elicited broad interpretations that overlapped a great deal with other phrases’ interpretation (particularly the phrases “fundamental”, “important”, and “valuable” on the significance/importance dimension (Figure 1) and “compelling”, “convincing”, and “useful” on the strength of support dimension (Figure 2). This suggests these phrases are not ideal for discriminating between studies with different degrees of importance and strength of support. If the same phrases often mean different things to different people, there is a danger of miscommunication between the journal and its readers. Responses on the significance/importance dimension were largely confined to the upper half of the scale, which is unsurprising, given the absence of negative phrases. It is unclear if the exclusion of negative phrases was a deliberate choice on the part of eLife’s leadership (because articles with little importance would not be expected to make it through editorial triage) or an oversight (see Footnote 1). Most participants’ implied rankings of the phrases were misaligned with the ranking intended by eLife —20% of participants had aligned rankings on the significance/importance dimension and 15% had aligned rankings on the strength of support dimension (Figure 3). The degree of mismatch was typically in the range of one or two discordant ranks (Figure 4). Heat maps (Figure 5) highlighted that phrases in the middle of scale (e.g., “solid”, “convincing”) were most likely to have discordant ranks.

By contrast, phrases in the alternative vocabulary tended to elicit more consistent interpretations across participants, and interpretations that had less overlap with other phrases (Figures 1 and 2; Tables 3 and 4). The alternative vocabulary was more likely to elicit implied rankings that matched the intended ranking — 62% of participants had aligned rankings on the significance/importance dimension and 67% had aligned rankings on the strength of support dimension (Figure 3). Mismatched rankings were usually misaligned by one rank (Figure 4). Although the alternative vocabulary had superior performance to the eLife vocabulary, it was nevertheless imperfect. Specifically, interpretation of phrases away from the middle of the scale on both dimensions (e.g., “low importance” and “very low importance”) tended to have some moderate overlap (Figures 1, 2 and 5). We do not know what caused this overlap, but, as discussed in the next paragraph, one possibility is that it is overly optimistic to expect peoples’ intuitions to align when they judge phrases in isolation, without any knowledge of the underlying scale.

Rather than presenting evaluative phrases in isolation (as occurs for eLife readers and occurred for participants in our study), informing people of the underlying ordinal scale may help to improve communication of evaluative judgements. eLife could refer readers to an external explanation of the vocabulary, however, prior research on interpretation of probabilistic phrases suggests this may be insufficient as most people neglect to look up the information (Budescu et al., 2014; Wintle et al., 2019). A more effective option might be to explicitly present the phrases in their intended ordinal structure (Wintle et al., 2019). For example, the full importance scale could be attached to each evaluation statement with the relevant phrase selected by reviewers/editors highlighted (Figure 6a). Additionally, phrases could be accompanied by mutually exclusive numerical ranges (Figure 6b); prior research suggests that this can improve consistency of interpretation for probabilistic phrases (Wintle et al., 2019). It is true that the limits of such ranges are arbitrary, and editors may be concerned that using numbers masks vague subjective evaluations in a veil of objectivity and precision. To some extent we share these concerns, however the goal here is not to develop an ‘objective’ measurement of research quality, but to have practical guidelines that improve accuracy of communication. Specifying a numerical range may help to calibrate the interpretations of evaluators and readers, so that the uncertainty can be accurately conveyed.

**Figure 6.**
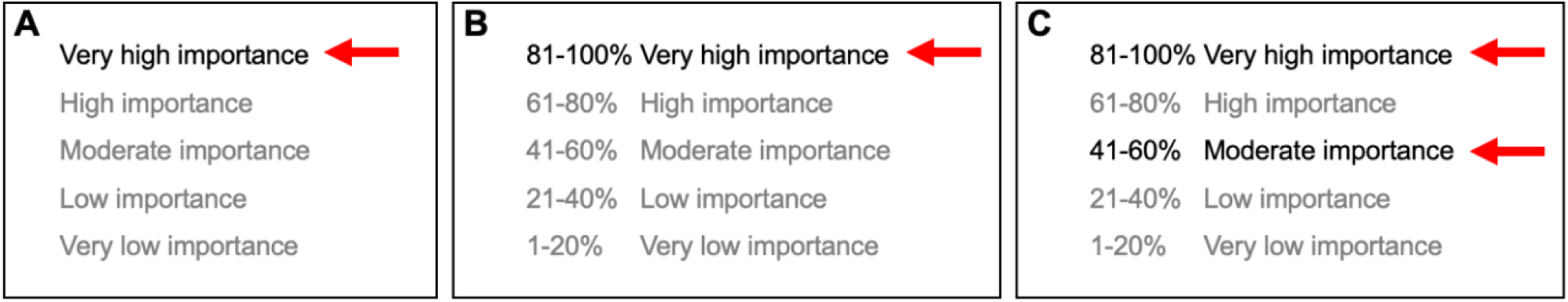
Explicit presentation of the intended ordinal structure (A-C), potentially with numerical ranges (B), could improve consistency of interpretation. Judgements by multiple reviewers could also be represented (by different arrows) without forcing consensus (C).

Our study has several important limitations. First, we did not address whether editor/reviewer opinions provide (a) valid assessments of studies; or (b) whether the vocabularies provide valid measurements of those opinions. We also note that eLife assessments are formed via consensus, rather than representing the opinions of individuals, which raises questions about how social dynamics may affect the evaluation outcomes. It may be more informative to solicit and report individual assessments from each peer reviewer and editor, rather than force a consensus (e.g., see Figure 6C). Although these are important issues, they are beyond the scope of this study, which is focused on clarity of communication.

Second, we are particularly interested in how the readership of eLife interpret the vocabularies, but because we do not have any demographic information about the readership, we do not know the extent to which our sample is similar to that population. We anticipated that the most relevant demographic characteristics were education status (because the content is technical), knowledge of subject area (because eLife publishes biomedical and life sciences), and language (because the content is in English). All of our participants reported speaking fluent English, the vast majority had doctoral degrees, and about one third had a degree in the Biomedical and Life Sciences. Relative to this sample, we expect the eLife readership probably consists of more professional scientists, but otherwise we think the sample is likely to be a good match to the target population. Also note that eLife explicitly states that eLife assessments are intended to be accessible to non-expert readers (*eLife’s New Model*, 2022), therefore, our sample is still a relevant audience, even if it might contain fewer professional scientists than eLife’s readership.

Third, to maintain experimental control, we presented participants with very short statements that differed only in terms of the phrases we wished to evaluate. In practice however, these phrases will be embedded in a paragraph of text (e.g., Box 1) which may also contain “aspects” of the vocabulary definitions (Table 1) “when appropriate” (*eLife’s New Model*, 2022). It is unclear if the inclusion of text from the intended phrase definitions will help to disambiguate the phrases and future research could explore this.

Fourth, participants were asked to respond to phrases with a point estimate, however, it is likely that a range of plausible values would more accurately reflect their interpretations (Reagan et al., 1989; Wallsten et al., 1986). Because asking participants to respond with a range (rather than a point estimate) creates technical and practical challenges in data collection and analysis, we opted to obtain point estimates only.

## Conclusion

Overall, our study suggests that using more structured and less ambiguous language can improve communication of research evaluations. Relative to the eLife vocabulary, participants’ interpretations of our alternative vocabulary were more likely to align with each other, and with the intended interpretation. Nevertheless, some phrases in the alternatively vocabulary were not always interpreted as we intended, possibly because participants were not completely aware of the vocabulary’s underlying ordinal scale. Future research, in addition to finding optimal words to evaluate research, could attempt to improve interpretation by finding optimal ways to present them.

## SUPPLEMENTARY INFORMATION A: Example stimuli and attention check

This section contains example stimuli and the attention check for illustrative purposes. The veridical stimuli are available on the Open Science Framework (https://osf.io/jpgxe/).

**Supplementary Figure A1.**
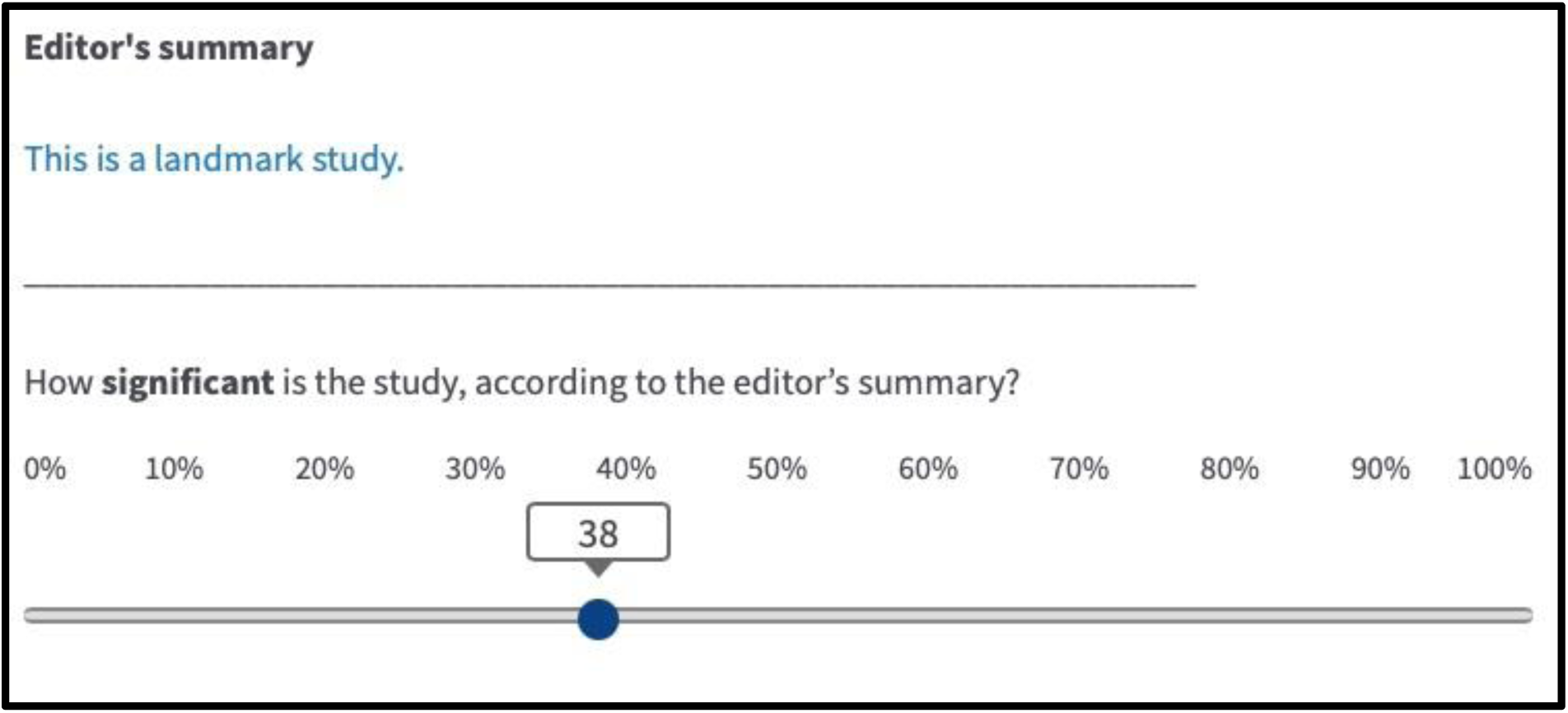
An example summary statement referring to a study’s significance and the corresponding response scale with an arbitrary response shown.

**Supplementary Figure A2.**
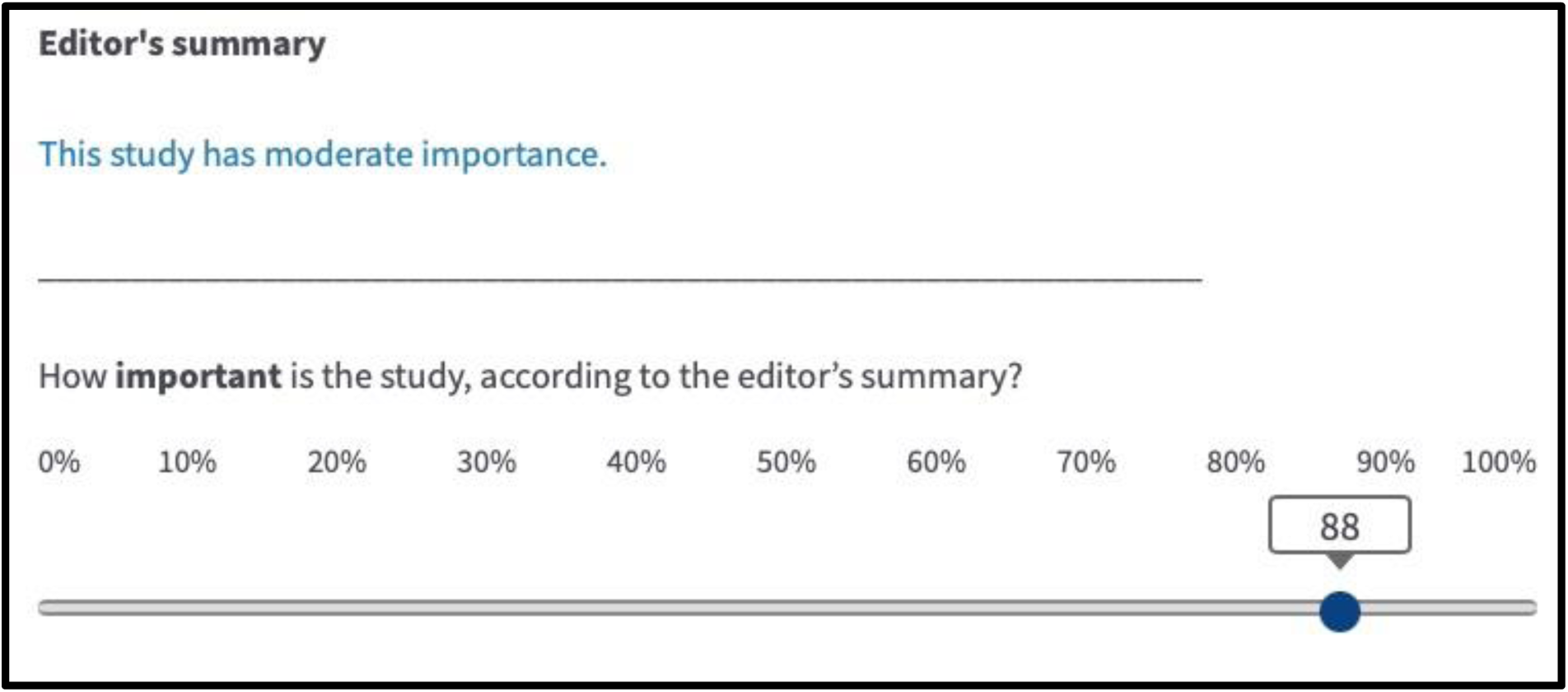
An example summary statement referring to a study’s importance and the corresponding response scale with an arbitrary response shown.

**Supplementary Figure A3.**
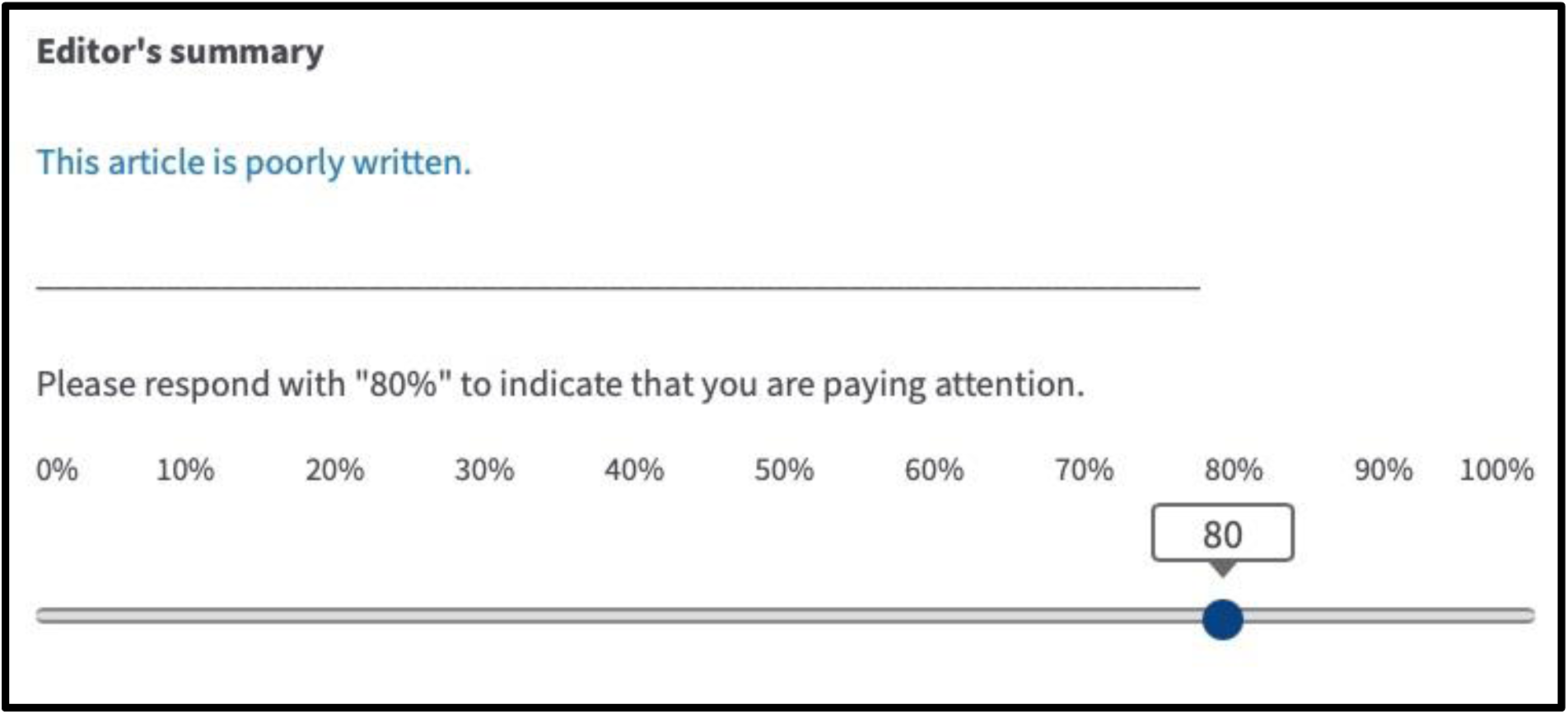
Attention check statement.

## SUPPLEMENTARY INFORMATION B: Sample size planning

We firstly decided that 300 participants was a reasonable sample size target given our resources. We then evaluated the expected statistical power and precision in a plausible scenario, assuming a sample size of 300 and a two-sided test with alpha = .05. For shorthand, we refer to a ‘correct response’ where the observed ranking matches the intended ranking. In a scenario where 10% of people respond correctly to the eLife vocabulary and incorrectly to the alternative vocabulary, and 20% of people respond correctly to the alternative vocabulary and incorrectly to the *eLife* vocabulary, this would yield a McNemar odds ratio of 2, 95% confidence intervals [1.3-3.1] and statistical power of 0.87 with an exact McNemar test. Analysis code documenting these calculations is available at https://osf.io/8v9n5 (under the heading “Sample size planning”).

## SUPPLEMENTARY INFORMATION C: Task instructions

This section contains copies of the task instructions for illustrative purposes. The veridical task instructions are available on the Open Science Framework (https://osf.io/jpgxe/).

Before starting the study, participants will be presented with the instructions shown in Supplementary Box C1 and have the opportunity to respond to a practice statement. At the start of each block they will be shown the instructions shown in Supplementary Box C2.

### Supplementary Box C1. Pre-study instructions

PAGE 1

Thank you for your participation in this study. Please ensure you are in a quiet, distraction-free environment before starting the task. Please give the task your full attention, it will only take about 10 minutes of your time.

PAGE 2

Imagine that you visit the website of a scientific journal to read some articles reporting scientific studies. You see that each article is accompanied by a short summary statement expressing the editor’s opinion of the study report in the article.

We are going to show you 22 statements describing the editor’s opinion of 22 different research studies. For each statement, we’d like you to tell us what you think about a particular aspect of the study using a slider on a scale ranging from 0 to 100%.

After each statement, you will complete a 15 second task involving simple multiplication questions.

PAGE 3

Here’s an example before we start. The editor’s summary appears in blue. In this example, your task is to rate how clearly written you think the article is based on the editor’s summary statement. You can click and drag the slider to choose your response.

Practice using the slider now and then click next when you are ready to start the study. If you are having technical problems or anything is unclear, please contact

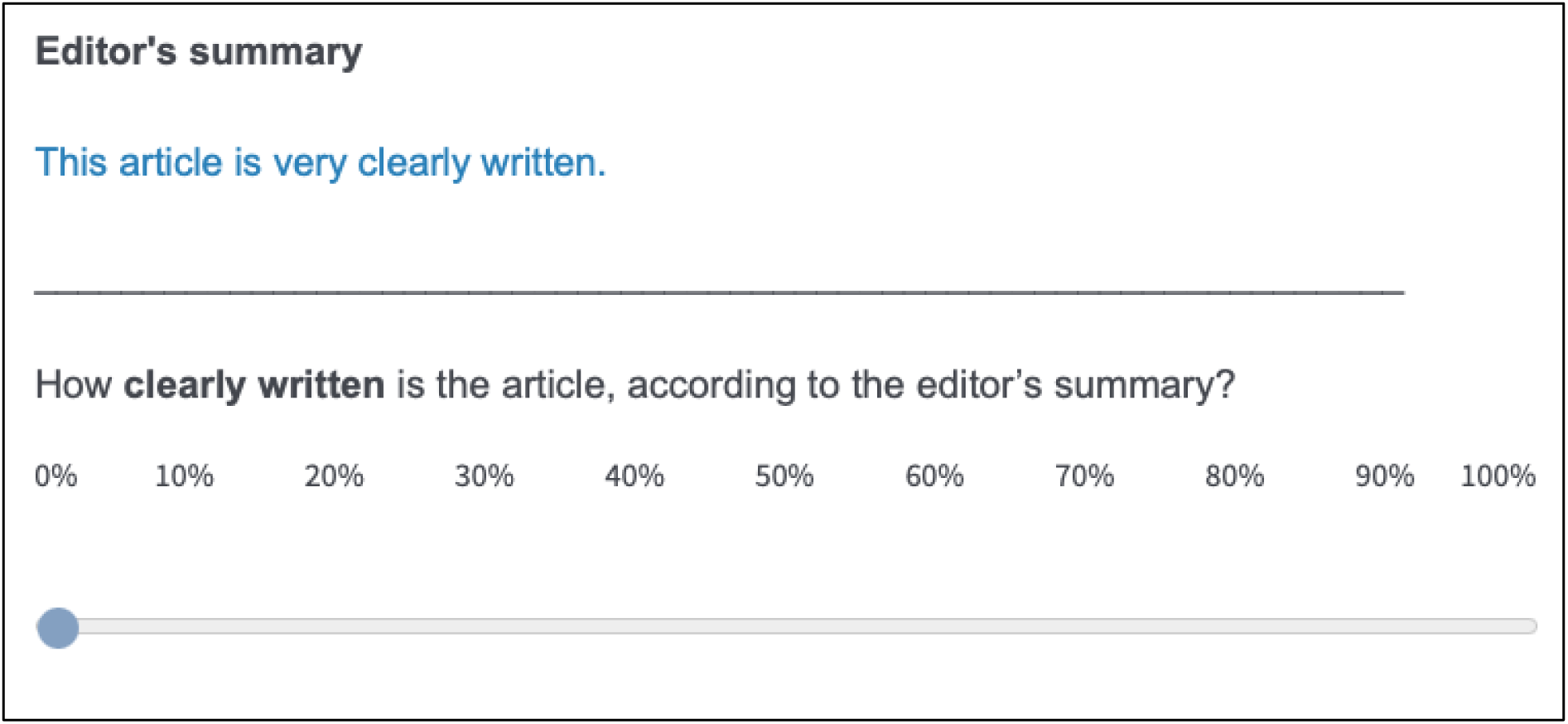

### Supplementary Box C2. Pre-block instructions

STRENGTH OF SUPPORT BLOCKS

You will now see statements about the strength of support offered by 5/6 different research studies. These statements represent the journal editor’s opinion about the strength of support each study offers towards its main claims.

SIGNIFICANCE BLOCK

You will now see statements about the significance of 5 different research studies. These statements represent the journal editor’s opinion about the significance of each of the study’s main claims.

IMPORTANCE BLOCK

You will now see statements about the importance of 5 different research studies. These statements represent the journal editor’s opinion about the importance of each of the study’s main claims.

## STAGE TWO REGISTERED REPORT

**SUPPLEMENTARY INFORMATION D.** Peer Community in Registered Reports Design Template.

This study was originally peer reviewed and received in-principle acceptance from Peer Community in Registered Reports. That platform required the Design Template below.

**Supplementary Table D1.**
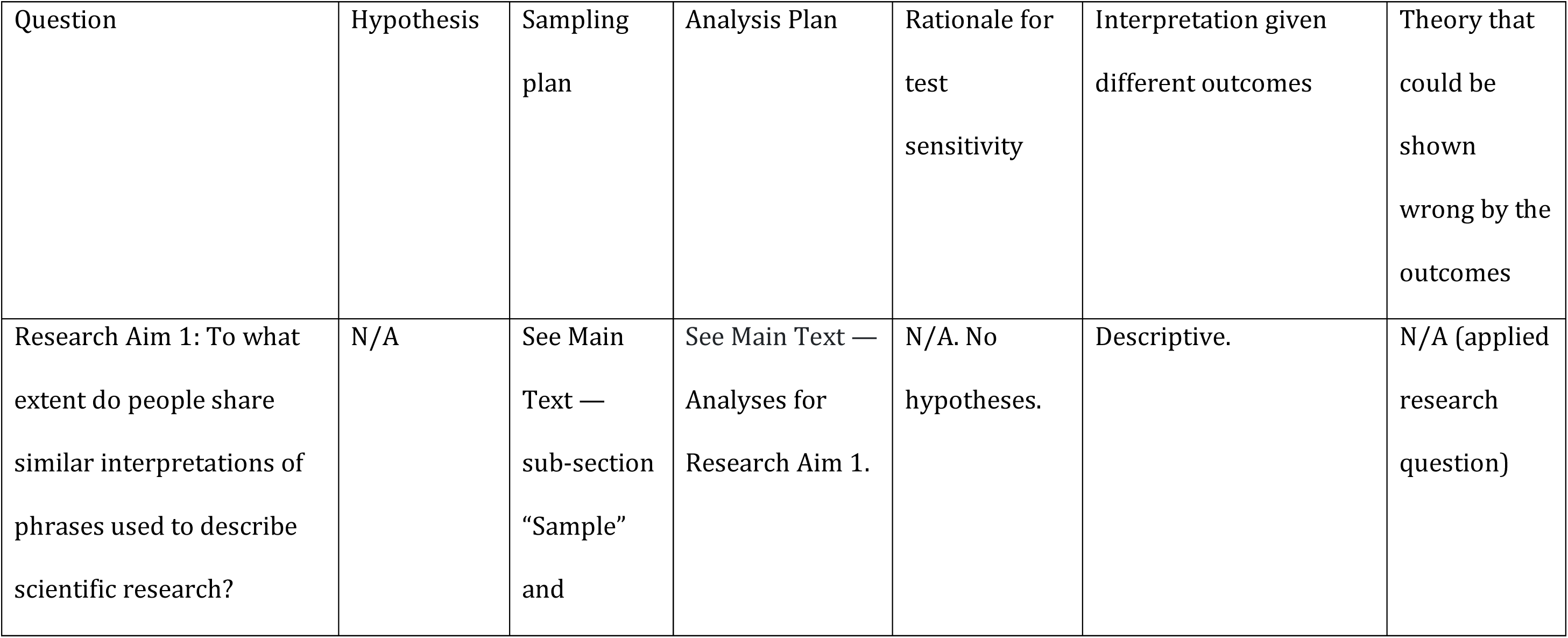
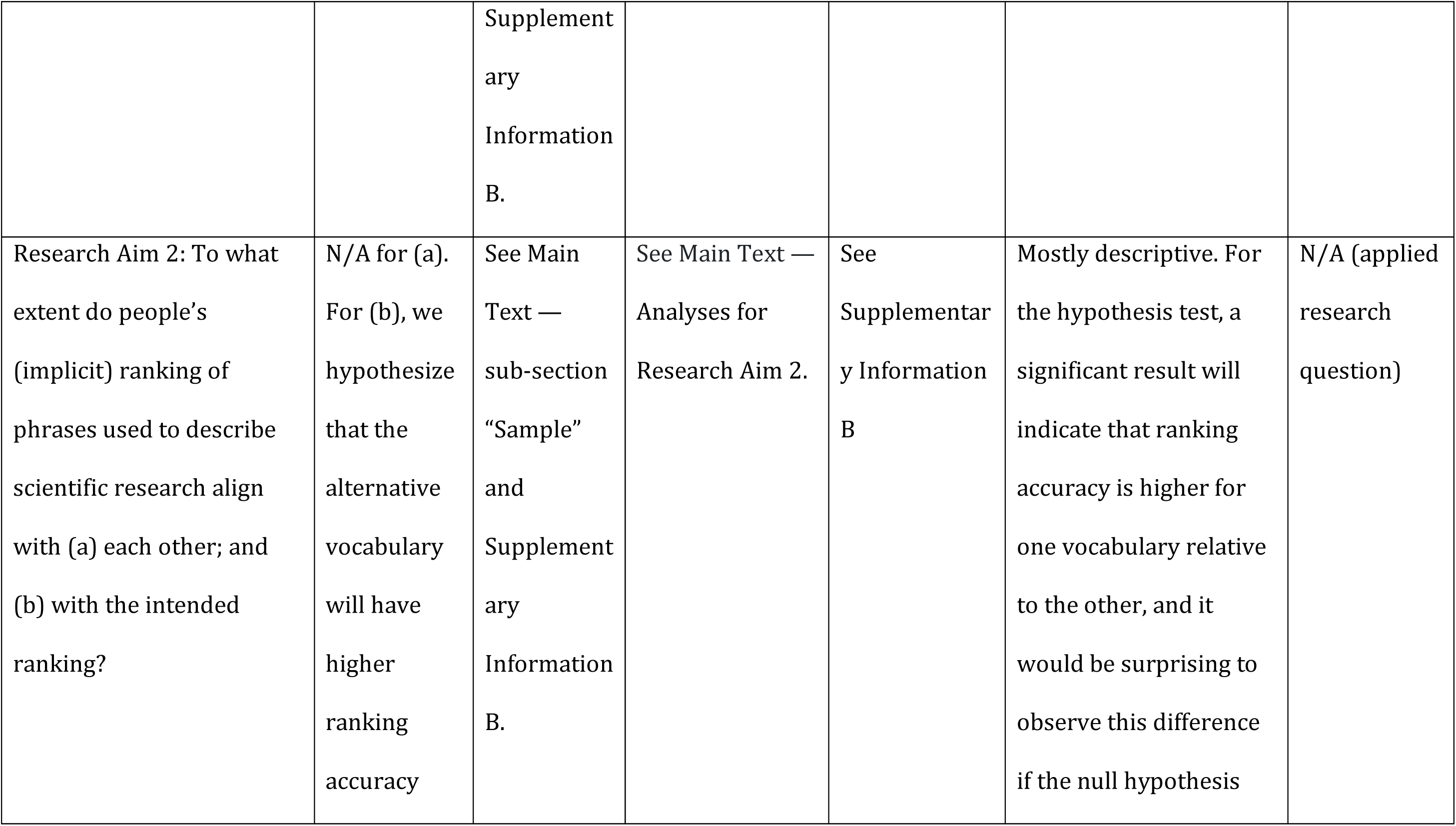
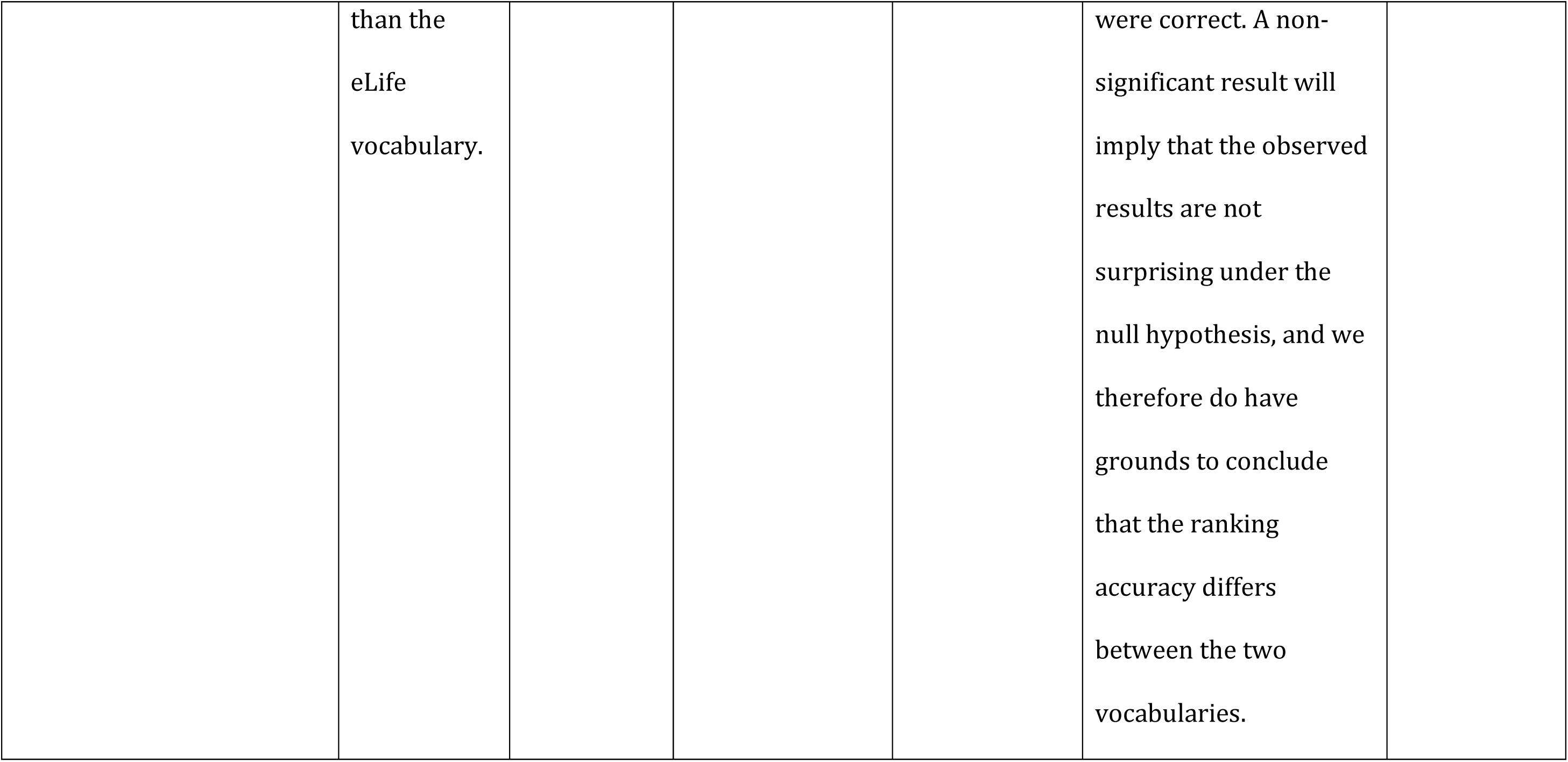
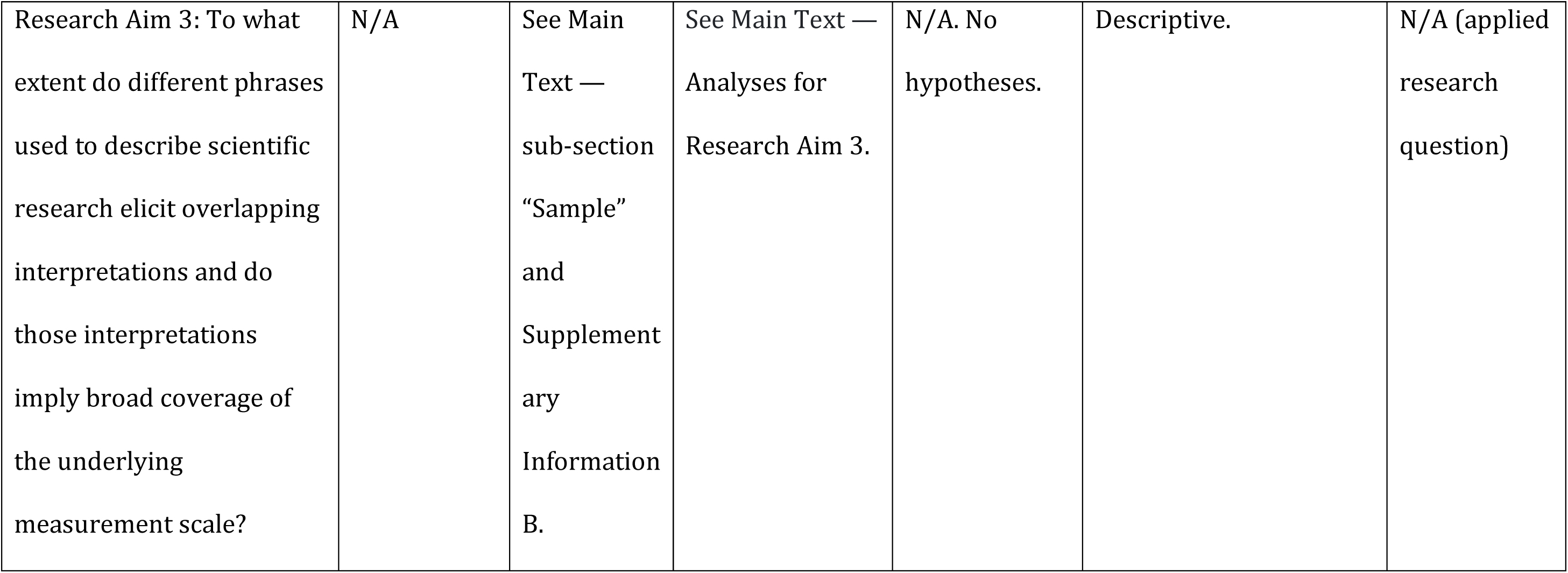
Peer Community in Registered Reports Design Template.

**SUPPLEMENTARY INFORMATION E.** Expanded versions of Table 3 and Table 4 percentile estimates with confidence intervals.

**Supplementary Table E1:**
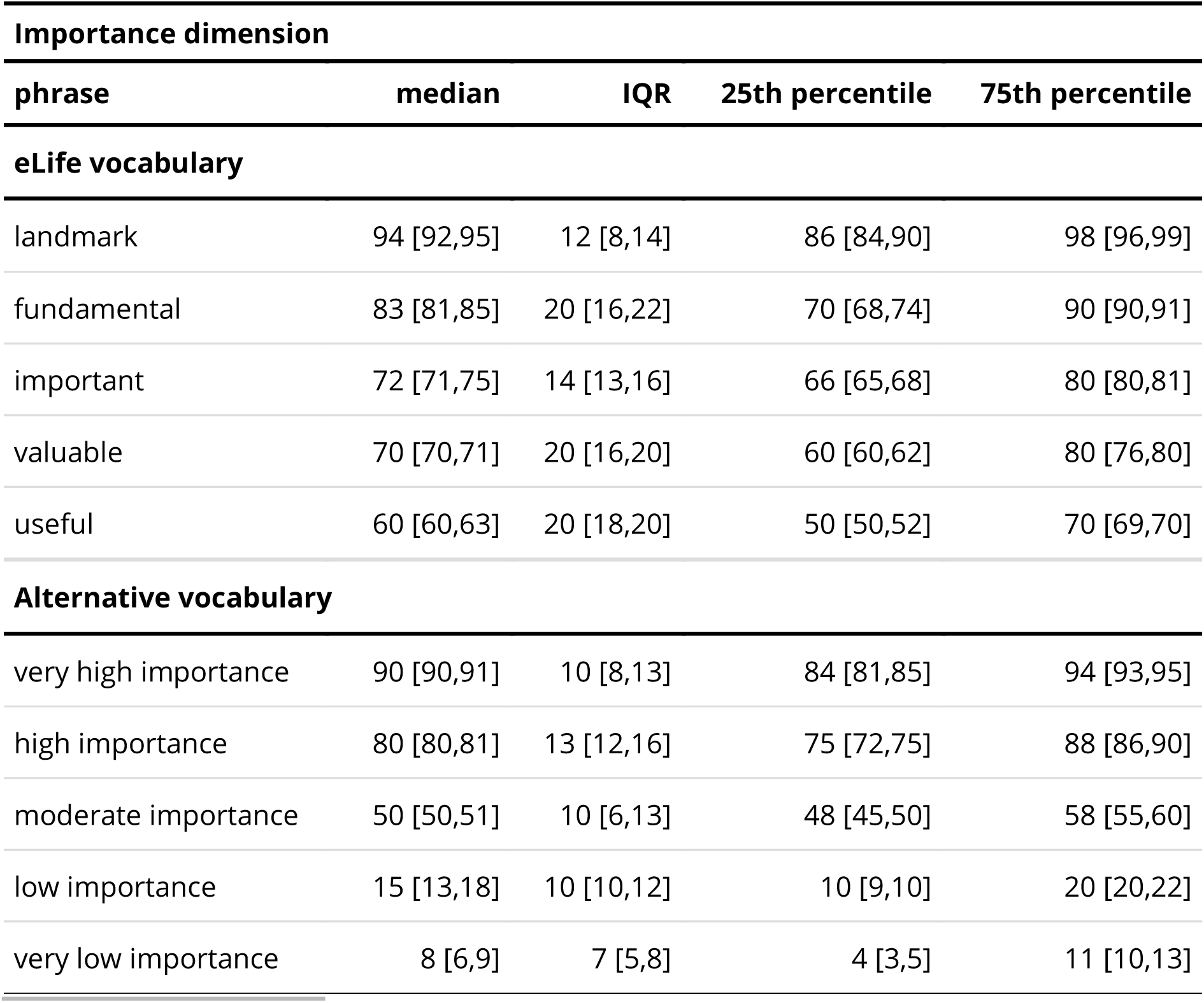
Percentile estimates for participant responses to phrases on the significance/importance dimension for eLife and alternative vocabularies. 95% confidence intervals bootstrapped with the percentile method are shown in the square brackets. IQR: Interquartile range.

**Supplementary Table E2:** Percentile estimates for participant responses to phrases on the support dimension for eLife and alternative vocabularies. 95% confidence intervals bootstrapped with the percentile method are shown in the square brackets. IQR: Interquartile range.

| **Support dimension** |  |  |  |  |
| --- | --- | --- | --- | --- |
| **phrase** | **median** | **IQR** | **25th percentile** | **75th percentile** |
| **eLife vocabulary** |  |  |  |  |
| exceptional | 95 [94,95] | 8 [7,10] | 90 [90,90] | 98 [97,100] |
| compelling | 80 [76,80] | 16 [14,19] | 70 [70,71] | 86 [85,90] |
| convincing | 75 [71,75] | 16 [14,20] | 65 [63,67] | 81 [80,85] |
| solid | 74 [71,75] | 17 [15,18] | 65 [64,67] | 82 [80,83] |
| incomplete | 20 [16,20] | 20 [19,22] | 10 [8,10] | 30 [28,30] |
| inadequate | 10 [10,13] | 15 [13,16] | 5 [4,5] | 20 [18,21] |
| **Alternative vocabulary** |  |  |  |  |
| very strong support | 88 [85,90] | 12 [10,14] | 80 [80,82] | 92 [91,94] |
| strong support | 77 [75,80] | 15 [11,15] | 70 [70,71] | 85 [81,85] |
| moderate support | 50 [50,51] | 10 [7,13] | 46 [45,49] | 56 [55,59] |
| weak support | 15 [14,18] | 12 [10,15] | 10 [8,10] | 22 [20,25] |
| very weak support | 9 [7,10] | 8 [5,10] | 5 [4,5] | 13 [10,14] |

**SUPPLEMENTARY INFORMATION F.** Frequency of implied rankings.

**Supplementary Table F1.** Implied rankings for the *eLife* vocabulary, significance/importance dimension.

| Implied ranking | n | % [CI] |
| --- | --- | --- |
| 1-2-3-4-5 | 59 | 20 [15, 24] |
| 1-3-2-4-5 | 32 | 11 [6, 15] |
| 1-2-3-5-4 | 20 | 7 [2, 11] |
| 2-1-3-4-5 | 19 | 6 [2, 11] |
| 1-4-3-2-5 | 15 | 5 [1, 10] |
| 1-3-2-5-4 | 14 | 5 [0, 9] |
| 1-2-4-3-5 | 13 | 4 [0, 9] |
| 2-3-4-1-5 | 13 | 4 [0, 9] |
| 1-4-2-3-5 | 10 | 3 [0, 8] |
| 2-4-3-1-5 | 9 | 3 [0, 8] |
| 3-2-1-4-5 | 9 | 3 [0, 8] |
| 2-3-1-4-5 | 8 | 3 [0, 7] |
| 2-1-4-3-5 | 6 | 2 [0, 7] |

**Table note:** 42 other rankings with $n < 6$ are not shown here. CI: Confidence interval.

**Supplementary Table F2.**
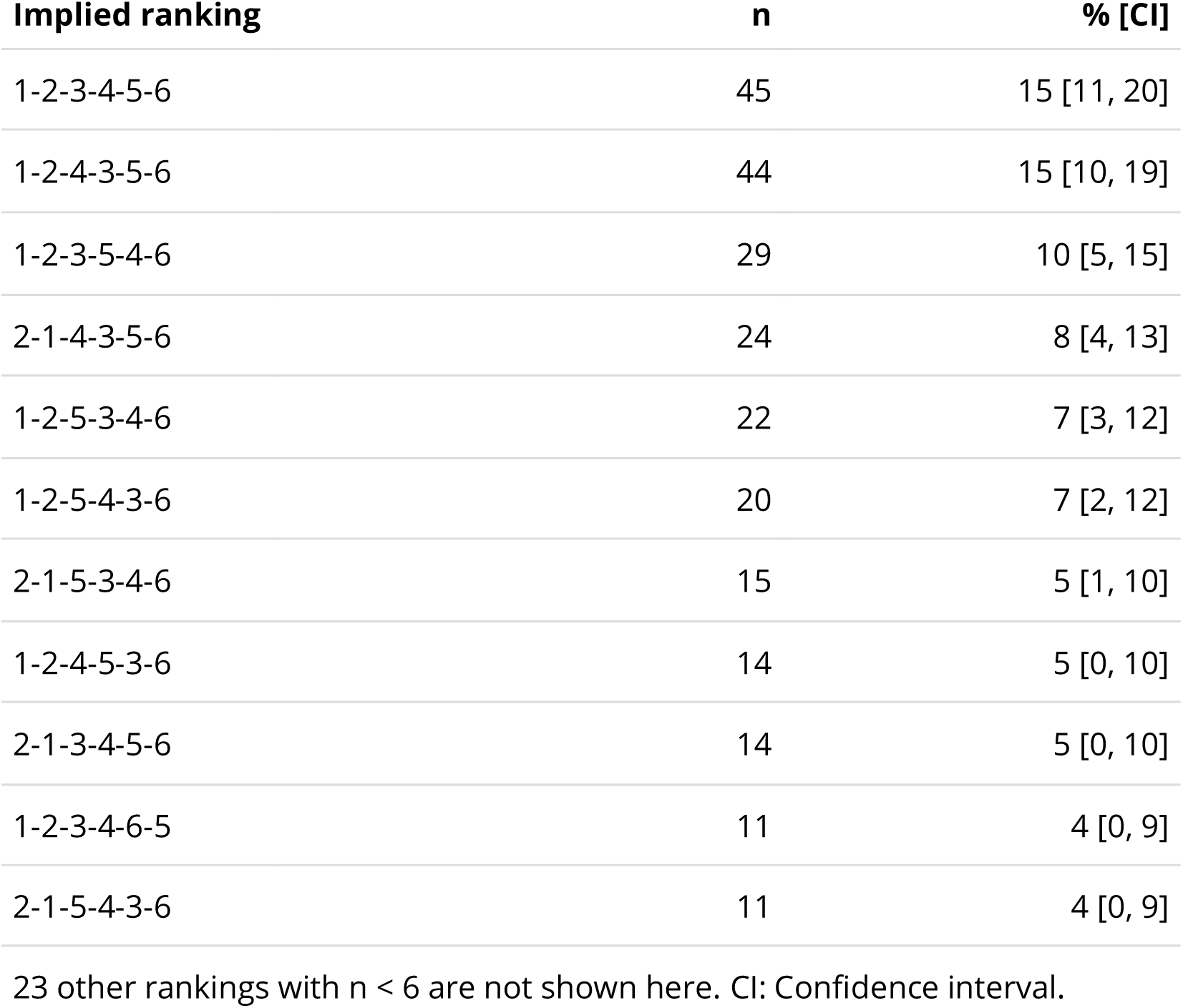
Implied rankings for the eLife vocabulary, support dimension.

| Implied ranking | n | % [CI] |
| --- | --- | --- |
| 1-2-3-4-5-6 | 45 | 15 [11, 20] |
| 1-2-4-3-5-6 | 44 | 15 [10, 19] |
| 1-2-3-5-4-6 | 29 | 10 [5, 15] |
| 2-1-4-3-5-6 | 24 | 8 [4, 13] |
| 1-2-5-3-4-6 | 22 | 7 [3, 12] |
| 1-2-5-4-3-6 | 20 | 7 [2, 12] |
| 2-1-5-3-4-6 | 15 | 5 [1, 10] |
| 1-2-4-5-3-6 | 14 | 5 [0, 10] |
| 2-1-3-4-5-6 | 14 | 5 [0, 10] |
| 1-2-3-4-6-5 | 11 | 4 [0, 9] |
| 2-1-5-4-3-6 | 11 | 4 [0, 9] |

23 other rankings with $n < 6$ are not shown here. CI: Confidence interval.

**Supplementary Table F3.** Implied rankings for the alternative vocabulary, significance/importance dimension.

| Implied ranking | n | % [CI] |
| --- | --- | --- |
| 1-2-3-4-5 | 188 | 62 [57, 68] |
| 2-1-3-4-5 | 50 | 17 [11, 22] |
| 1-2-3-5-4 | 35 | 12 [6, 17] |
| 2-1-3-5-4 | 23 | 8 [2, 13] |

Table note: 4 other rankings with $n < 6$ are not shown here. CI: Confidence interval.

**Supplementary Table F4.** Implied rankings for the alternative vocabulary, significance/importance dimension.

| Implied ranking | n | % [CI] |
| --- | --- | --- |
| 1-2-3-4-5 | 201 | 67 [62, 72] |
| 2-1-3-4-5 | 49 | 16 [11, 22] |
| 1-2-3-5-4 | 24 | 8 [3, 13] |
| 2-1-3-5-4 | 20 | 7 [2, 12] |

Table note: 6 other rankings with $n < 6$ are not shown here. CI: Confidence interval.

**SUPPLEMENTARY INFORMATION G. Further explanation of Kendall’s distance.**

Consider an example case where two individuals are asked to rank three fruits in their favourite order and we want to compare the similarity of the rankings. The individuals provide the following rankings:

*Person A = Orange, Apple, Pear*

*Person B = Pear, Apple, Orange*

In this case, Kendall’s distance (*Kd*) = 3, because three adjacent pairwise swaps are required to convert Person A’s list into Person B’s list (specifically, we need to swap Apple-Pear, then Pear-Orange, then Apple-Orange.). In this example the maximum *Kd* is 3, so these two rankings are maximally dissimilar.

For our study, we can use *Kd* the measure the similarity between a given participant’s ranking and the intended ranking. For example, we could compare:

*Intended (eLife) ranking = useful, valuable, important, fundamental, landmark (A hypothetical) observed ranking = useful, important, valuable, fundamental, landmark*

In this case, *Kd* = 1, because only a single adjacent pairwise swap (valuable <-> important) is necessary to convert the participant’s ranking into the intended ranking.

**SUPPLEMENTARY INFORMATION H. Demographics of Prolific members**

Complete demographic information about Prolific members is not available as demographic screening questions are voluntary. Based on the available responses at the time of our study, 30% of Prolific members said they were aged between 18-25, 58% said they were aged between 26-50, and 12% said they were aged between 51-100; asked about their gender, 35% identified as a man, 46% identified as a woman, and 2% identified as non-binary; 18% said they were currently a student; 87% said they were fluent English speakers; 31% said they have UK nationality and 31% said they have USA nationality; when asked to report the highest level of education they have completed, 25% said undergraduate degree, 12% said graduate degree, and 2% responded doctorate degree.

## Footnotes

1 Note that eLife does not intend to send all submitted manuscripts for peer review (*eLife Review Process FAQs*, 2023), so it is possible they have omitted negative phrases on the grounds that less-than-useful papers should not make it through initial editorial triage. Nevertheless, we think it would be prudent to include negative phrases in the vocabulary because peer reviewers may disagree with the editor, or the editor may change their mind.

2 We suggest that if eLife wants to convey breadth/scope as well as degree, this should be represented by a separate dimension.

3 All participants had previously responded to a Prolific pre-screening question that their highest completed education level was a doctoral degree.

## Notes

**Funding statement:** This study was supported by funding from the Melbourne School of Psychological Sciences, University of Melbourne.

### Competing Interest Statement

Simine Vazire is a member of the PLOS Board of Directors. All other authors declare they have no conflicts of interest.

